# Bouncing back: Transient, but not long-lasting, effects of background disturbances on speech processing

**DOI:** 10.64898/2026.07.13.738235

**Authors:** Orel Levy, Rotem Waxman, Shun Okada, Ofek Ben-Abu, Elana Zion-Golumbic

**Affiliations:** The Gonda Center for Brain Research, Bar Ilan University

## Abstract

Processing and understanding speech in face of occasional background disturbances can be difficult and often leads to distraction and reduced comprehension. However, despite the vast literature on the detrimental effects of processing speech-in-noise, little is known about the time-course of these effects. Here we studied whether reduced speech processing is limited only to the period when disruptions mask target speech, or if processing difficulties linger even after disruptions end, suggesting longer lasting distraction. Using an ecologically-relevant experimental design, we measured neural activity and skin-conductance from participants as they watched an educational video-lecture. Short background speech comments, mimicking plausible classroom disruptions, were occasional embedded in half of the lecture segments. Speech disruptions elicited robust neurophysiological orienting responses, reflected in a neural auditory ERP and a phasic increase in skin conductance. Importantly, when disruptions were presented, we observed a marked reduction in neural tracking of the lecture, increased inter-subject correlation (ISC), and decreased alpha power, suggesting a brief diversion of processing resources away from the target speech. However, these effects were transient in nature, with all neural measures returning to baseline immediately after disruption offset, showing no evidence for prolonged disruption-related costs or impaired comprehension. Interestingly, neurophysiological responses to disruption and return to baseline were not correlated with self-reported symptoms of attention-deficits (ADD). These findings highlight the resilience of auditory attention, demonstrating its ability to maintain a robust representations of target speech over time, despite of transient ecological disturbances.

## Introduction

Understanding speech processing in dynamic real-life environments is a core translational challenge of cognitive neuroscience. Perhaps the most challenging aspect of real-life speech processing is dealing with background noises and disturbances, which can both mask target speech (Brungart, 2001; Mattys et al., 2012; Wang & Xu, 2021) and distract attention away from it (Bell et al., 2022). The detrimental effects of background noise on speech processing are well-documented, as manifest in reduced speech intelligibility, comprehension and memory for speech content (Klatte et al., 2013; Mattys et al., 2012; Socher et al., 2026), and are accompanied by poorer neural representation of target speech (Broderick et al., 2018; Dai et al., 2022; Ding & Simon, 2013; Etard & Reichenbach, 2019; Levy, Korisky, et al., 2025). And yet, not all types of noise are disruptive to the same degree. For example, intelligible speech and semantically-meaningful sounds are often more disruptive than meaningless sounds or white noise, indicating that the competition for processing resources between concurrent stimuli extends beyond low-level acoustic masking (Brungart, 2001; Dai et al., 2022; Lecumberri & Cooke, 2006; Schneider et al., 2007; Tun et al., 2002; Van Engen & Bradlow, 2007). Similarly, structured noise with frequent onsets can be more disruptive than continuous noise, suggesting that the temporal structure of competing sounds plays a role in a listener’s ability to suppress them (Bacon & Viemeister, 1985; Levy, Korisky, et al., 2025; Levy, Shadi, et al., 2025; Wier et al., 1977). Specifically, it has been proposed that background noise that contains transient salient events, may induce repeated re-sets to the auditory system, preventing adaptation and ultimately leading to more severe disruption (Anstis & Saida, 1985; Bregman, 1978; Korte et al., 2025; McFadden et al., 2010; Rankin et al., 2017). However, since most studies of speech-in-noise use designs where utterances are continuously masked, little is known about the effects of transient masking events on speech processing and their time course of disruption.

The auditory system is highly sensitive to local changes in the acoustic scene, with unexpected events generating heightened sensory responses (such as the N100 and Mismatch Negativity; MMN) (Näätänen & Alho, 1995; Näätänen & Picton, 1987; Sams et al., 1991; Vaughan & Ritter, 1970), and eliciting later surprise-related responses associated with involuntary attentional capture or increased arousal (between 250-350 ms, referred to sometimes as the Novelty-P3) (Baragona et al., 2025; Fiedler et al., 2025; Friedman et al., 2001). In addition, unexpected sounds are accompanied by increased physiological arousal, measured through skin conductance (Koelsch et al., 2008; Widmann et al., 2018) and pupil dilation (Baragona et al., 2025; Smith et al., 2025). Although, these effects have mostly been demonstrated using artificial paradigms and simple stimuli, such as the well-known oddball task (Justen & Herbert, 2018; Kim, 2014; Näätänen et al., 2004; Squires et al., 1975), they do seem to generalize to more ecological contexts, including environmental sounds, human-made non-speech sounds, personally relevant words and semantic incongruencies (Bidet-Caulet et al., 2015; Brown et al., 2023; Fiedler et al., 2025; Huang & Elhilali, 2020; Levy, Libman Hackmon, et al., 2025; Nguyen et al., 2024; Stavropoulos & Carver, 2016). Given that transient events elicit strong, and to some degree automatic, neural and physiological responses, this raises an important question: How does hearing semantically salient yet task-irrelevant speech, affect ones’ ability to maintain attention to- and comprehend target speech?

This question addresses a tension that is fundamental to attentional functioning in general, and attention to speech in particular: how to balance the allocation of processing resources between target and task-irrelevant stimuli. On the one hand, the essence of selective attention is to amplify target stimuli and suppress or “filter out” task-irrelevant stimuli to prevent distraction, a top-down process that can be effortful but is critical for optimal task performance (Baluch & Itti, 2011; Chait et al., 2010; Kaufman & Zion Golumbic, 2023; Zion Golumbic et al., 2013). On the other hand, our sensory systems must retain bottom-up sensitivity and respond to unexpected peripheral events, and re-orient towards them if needed (Buschman & Miller, 2007; Corbetta & Shulman, 2002; Huang & Elhilali, 2020; Kaya & Elhilali, 2014). For irrelevant stimuli that are transient in nature, this tension can be addressed at two different time points: How is speech processing affected **during** transient disruptions? And what happens **after** the disruption ends **-** does the system recover rapidly or are there lingering effects on speech processing? To date, the few studies investigating the effects of transient disruptions on speech processing in a time-resolved manner have found reduced neural tracking of target speech, reduced gamma power and increased arousal when disruptions are presented, consistent with the idea of competition for cognitive resources and increased investment of effort (Fiedler et al., 2025; Huang & Elhilali, 2020; Straetmans et al., 2022). However, mixed results are reported regarding how long these effects last, with some findings indicating lingering suppression of target speech and increased arousal for several seconds (Fiedler et al., 2025; Straetmans et al., 2022), while others show evidence of rapid recovery (or even a rebound) of attention (Holtze et al. 2021).

The limited, and somewhat inconclusive, nature of current research, motivated us to study the effects of background disruptions on speech processing in an ecological context where it is particularly pertinent: watching a video-based lecture. Participants viewed a 20- minute educational lecture (the **target** stream) while we recorded their brain activity using EEG and their arousal via galvanic skin response (GSR). The lecture was divided into 90-second long segments, half of which contained occasional background disruptions. We chose to use speech-based disruptions, in the form of short comments (e.g., “I’m so tired”, “that is interesting”), which carry semantic and socially-relevant content alongside their acoustic masking, factors that potentially increase their power to draw attention. Neural activity (EEG) and skin-conductance (GSR) were analyzed to extract indices associated with stimulus processing, attention and arousal: including speech tracking of the lecture (Broderick et al., 2018; Kaufman & Zion Golumbic, 2023), event-related responses, and oscillatory power in the alpha band (often linked to attention and engagement; Clements et al., 2023; Magosso et al., 2019), alongside behavioral performance (e.g. answering comprehension questions about the lecture). Crucially, analyses were performed in a time-resolved manner, allowing us to separately evaluate effects during and after the presentation of disturbances. By using spoken material that simulate the challenge of dealing with verbal disruptions in real-life listening contexts, our work extends traditional attention research in more ecologically valid directions. Specifically, it aims to shed light on how the brain balances competing bottom-up and top-down demands in real time and evaluate the cost of transient disruptions to speech-processing and the time-course for re-engaging with target speech after disruptions.

## Methods

### Participants

Data were collected from 60 adult volunteers (33 females, 27 males), aged 18 to 52 years (mean age: 26.35 ± 6.87). Sample size was determined based on previous studies from our group assessing speech-tracking (Levy et al. 2025a,b), and is sufficient for detecting a between-condition effect size of d = 0.3 with power of 0.8 (G*power simulation). All participants were fluent Hebrew speakers, with self-reported normal hearing and no history of neurological disorders, besides ADHD. Five participants reported having a prior clinical diagnosis of ADHD and provided certified documentation. In addition, prior to the experiment, all participants completed the Adult ADHD Self-Report Scale (ASRS), a validated 18-item questionnaire assessing AD(H)D symptoms (ASRS-v1.1; Adler et al., 2006; Hebrew version: Zohar & Konfortes, 2010). The questionnaire was administered on-line using the Qualtrics platform ($ref). ASRS scores were calculated as the sum across all items, with a sum over 51 considered to be a “candidate for ADHD”. One participant was excluded from the GSR analysis due to missing physiological recording. Compensation was provided in the form of monetary payment or course credit. The study received approval from the Institutional Review Board at Bar-Ilan University, and all participants provided informed consent prior to the experiment. The experiment described here was collected as part of a larger study, that included multiple attention tasks over two sessions, as reported elsewhere (Waxman et al., 2026).

### Experimental Procedure

The experiment was programmed and presented to participants using OpenSesame software (Mathôt et al., 2012; https://osdoc.cogsci.nl). Participants were seated in a sound-attenuated room and viewed a ∼20 minute long video-recording of a lecture, entitled "The Extraordinary Brain", on the topic of Savant syndrome. The video was presented on a computer monitor in front of the participant and audio was played from a loudspeaker placed behind the monitor (95 cm from participant). The lecture was divided into 12 segments, lasting 98.7 ± 16.8 seconds each, that were presented sequentially as individual trials. After each segment, participants answered three multiple-choice questions about its content.

In half of the trials (“Disruption” trials), additional short speech-utterances were occasionally presented from a loudspeaker positioned to the rear-left of the participant (95 cm from participant), serving as disruptions. Disruptions were designed to simulate a fellow student sitting behind the participant and commenting on the situation, and consisted of short utterances, recorded in a young-adult female voice (e.g., “this is so interesting”, “it might rain”, “I’m so tired”; duration 2.22 ± 0.43 seconds). All utterances were prefaced with the Hebrew interjection “Wai” (which is akin to “Aw” or “Oh” in English), which was recorded separately and added to the beginning of each utterance. This ensured an acoustically identical and easily identifiable onset for all disruptions, while maintaining ecological relevance. Each Disruption trial included six such disruption events, separated by an inter-stimulus interval of 11.55 ± 0.49 seconds (range: 10.75–12.33s), resulting in a total of 36 disruption events across the experiment. Disruptions were equated for loudness and were presented at a slightly louder loudness level relative to the lecture (3.5 dB). The mapping of lecture segments to condition (Disruption vs. Quiet) was counterbalanced across participants, to avoid material-specific effects. Comprehension accuracy was computed for each participant as the percentage of correct responses on the multiple-choice questions, averaged separately for the two conditions (Quiet, Disruptions). All participants had accuracy levels well above the 33% chance level, indicating task-compliance. Accuracy scores were compared between conditions using a two-tailed paired t-test.

### EEG and GSR data acquisition

EEG data were recorded using a 64-channel BioSemi ActiveTwo system (sampling rate: 2048 Hz) with Ag-AgCl electrodes. Two external electrodes were placed on the mastoids for reference. Electrooculographic (EOG) signals were simultaneously measured by 4 additional electrodes, located above and below the right eye and on the external side of both eyes. The GSR was recorded using two passive Nihon Kohden electrodes placed on the fingertips of the index and middle fingers of participants’ non-dominant hand, also recorded through the BioSemi system at the same sampling rate. Room audio was recorded using an external microphone (Boya BY-CM5 USB condenser microphone, 44.1 kHz). All data streams (EEG, GSR, audio and digital triggers) were recorded synchronously via the Lab Streaming Layer for synchronization (LSL; Kothe et al., 2025, https://github.com/sccn/labstreaminglayer).

### EEG and GSR preprocessing

EEG data were preprocessed using MATLAB together with the FieldTrip toolbox (Oostenveld et al., 2011; version: 20220729; https://www.fieldtriptoolbox.org), and custom-written scripts. The raw data were initially re-referenced to the linked mastoids, detrended, and demeaned. A zero-phase, two-pass Butterworth IIR filter was then applied between 0.5 and 40 Hz to attenuate low-frequency drifts and high-frequency noise. Independent component analysis (ICA) was performed (using the Fieldtrip ft_componentanalysis function, method = ‘runica’) to identify and remove components associated with ocular movements and heartbeats, based on visual inspection (mean ± SD: 4.32 ± 1.30 components per participant). EOG channels were included in the decomposition to improve the identification of ocular artifacts, and excessively noisy electrodes were temporarily excluded from the ICA decomposition to enhance stability. Following ICA, these temporarily-excluded channels were reinstated and interpolated using spatial neighbors (neighbourdist = 0.15), either across all trials or on a per-trial basis depending on the pattern of noise (mean ± SD: 2.12 ± 1.75 channels per participant). Finally, data were visually inspected; deflections exceeding ±50 μV (non-ocular) were flagged as artifacts and removed from event-related analyses (see below). GSR data were analyzed using the Ledalab toolbox (Benedek & Kaernbach, 2010b) and custom-written MATLAB scripts. Data low-pass filtered using a 4th-order Chebyshev Type I filter (cutoff 16 Hz, 1 dB ripple), converted to nanoSiemens (nS), and downsampled to 16 Hz. Signals were visually inspected for artifacts and corrected via linear interpolation, before applying a Continuous Decomposition Analysis (CDA) to decompose the signal into tonic and phasic driver components.

### EEG and GSR analysis

The effects of disruptions on neural responses (EEG) and physiological responses (GSR) were examined at three levels: (1) Event-related responses to disruption onsets (“Wai”), (2) Global effects of disruptions on lecture processing, assessed at the full-trial level, and (3) Local effects of disruptions on lecture processing, *during* and immediately *after* the disruption.

#### (1) Event-related responses to disruption onsets (“Wai”)

EEG and GSR data were segmented relative to the onset to of disruptions, which all began with an identical word (“Wai”), to capture transient neural and skin-conductance responses to these events. Neural Event-Related Potentials (ERPs) to disruptions were estimated by epoching the EEG data between –100 ms to 700 ms relative to disruption onset, and applying baseline-correction (between –100 to 0 ms) and a low-pass filter (20 Hz, zero-phase). Trials containing artifacts identified during preprocessing were rejected (mean exclusion: 1.75 ± 2.7 trials per participant), and the remaining artifact-free epochs were averaged to extract ERPs for each participant. Event-related changes in skin-conductance following disruptions, reflecting autonomic responses, were evaluated using the phasic driver of the skin conductance response extracted from the GSR signal. In line with the slow time-scale of autonomic responses, the phasic driver was segmented between 0-8 seconds after disruption onset, baseline-corrected relative to the average signal between 0-1 seconds (peak response is typically between 3-5 sec), and averaged across trials to extract the transient skin-conductance response for each participant. These responses are reported primarily for descriptive purposes, as the experiment design did not include relevant between-condition- comparisons for these measures.

#### (2) Global Effects of disruptions – assessed at the full trial level

To examine whether disruptions influenced neural processing of the lecture, continuous EEG recordings were segmented to separate between trials containing disruptions and those without disruptions (Quiet). We analyzed three neural metrics and two skin-conductance metrics at the full-trial level, and compared them between conditions: a) neural speech tracking of the lecture, b) inter-subject correlation (ISC), c) alpha power, d) phasic and tonic skin-conductance.

##### a) Speech tracking analysis

*We* used linear temporal response function (TRF) models (mTRF Toolbox; Crosse et al., 2016) to quantify the linear relationship between the speech envelope (S) and the corresponding EEG activity (R). Speech envelope was extracted from the speech- audio using an equally spaced filterbank between 100 and 10,000 Hz, based on Liberman’s cochlear frequency map. Each narrowband signal was transformed using the Hilbert transform, converted to its absolute value, and summed across frequencies to generate the final broadband envelope. Envelopes were then min-max normalized to ensure comparable scaling. The neural data (R) consisted of the EEG signal, bandpass filtered between 0.8 and 20 Hz and normalized using z-scores. Both the stimulus envelope and the EEG signals were aligned in time and downsampled to 100 Hz to improve computational efficiency. Both encoding and decoding approaches were used to estimate speech-tracking responses. In the encoding model, the speech envelope was used to predict the EEG response (S → R; lags −150 to 450 ms relative to the stimulus), whereas in the decoding model the EEG signal was used to reconstruct the speech envelope (R → S; lags 0 to 400 ms). Model performance was quantified as the Pearson correlation between the predicted and the observed signals, referred to here as predictive power (encoding) and reconstruction accuracy (decoding). Multivariate models were used in both encoding and decoding, estimated separately for Disruption and Quiet trials. For Disruption trials, the model included two regressors: one with the envelope of the lecture-speech envelope and one with the envelope of disruptions. For the Quiet trials, one regressor contained the envelope of the lecture speech and a second regressor was set to zeros. This approach maintains symmetric model architecture for both conditions, while accounting for the fact that only the Disruption condition contained two concurrent stimuli. Encoding and decoding procedures used ridge-regression optimization, with the ridge regularization parameter (λ) estimated using a leave-one-out cross-validation procedure, to avoid overfitting. For the encoding model, λ values ranging from 10⁻² to 10⁵ were tested, and λ = 10 was selected for the entire group, which showed the highest average predictive power across participants and channels. As discussed previously in Har-Shai Yahav et al., 2024; Kaufman & Zion Golumbic, 2023, using a common λ-value across participants for encoding allows for more interpretable group-level TRF time-courses. For the decoding analysis, λ values ranging from 10⁻¹ to 10⁵ were tested, and optimized for each participant, selecting the λ value with maximal reconstruction accuracy.

Model significance was assessed using a permutation test, where null distributions were generated by shuffling the pairing between the stimulus (S) and neural response (R) across trials (100 permutations per participant). To avoid biasing, this analysis combined trials from both conditions. For encoding models, we identified electrodes whose predictive power exceeded the top 95^th^ percentile of the null distribution, and these were selected for use in subsequent encoding analyses (total: 40 electrodes; Figure 3A). We then tested for differences in predictive power and TRF time-course between Disruption and Quiet conditions, within these electrodes (paired t-tests, Bonferroni/cluster corrected for multiple comparisons). For decoding models, we established the reliability of the model by testing if real reconstruction accuracy fell with the top 95^th^ percentile of the null distribution. We then compared reconstruction accuracy values between Disruption and Quiet conditions using a paired t-test.

##### b) Inter subject correlation (ISC)

Inter-subject correlation (ISC) was used to quantify the similarity in ongoing neural activity across participants during the lecture, and to assess whether this shared neural activity was affected by the presence of Disruptions. Because ISC analysis requires that all participants are exposed to identical stimuli, this analysis was conducted separately for the two subgroups of participants who shared the same trial structure (in the counterbalanced design), and results were then averages across subgroups. ISC was computed using the publicly available Parra Lab code for EEG inter-subject correlation (Cohen & Parra, 2016; Ki et al., 2016; https://www.parralab.org/isc/). Specifically, correlated component analysis (CCA; Dmochowski et al., 2012) was applied to identify spatial components that maximized across-subject correlation. The first three components, which captured the strongest shared variance, were retained for analysis. To estimate the ISC for each participant, which reflects their similarity to the “group”, we calculated the correlation between the time-course of each component in single-participants data vs. the average time course of all other participants, and then averaged the correlation coefficients across the first three components to obtain a single measure per participant. ISC was computed separately for Disruption and Quiet trials condition, and differences between conditions were assessed using paired-sample t-tests.

##### c) Alpha power Analysis

*We* analyzed the power spectral density (PSD) of the continuous EEG signal to investigate whether the presence of Disruptions affects ongoing oscillatory activity. PSD was estimated using a multitaper Fast Fourier Transform (FFT) with Hanning tapers (2– 40 Hz), implemented in the FieldTrip toolbox. To separate oscillatory activity from the underlying aperiodic 1/f background, we applied the Fitting Oscillations and One-Over-F (FOOOF) algorithm (Donoghue et al., 2020). The analysis focused on the alpha frequency band (7–13 Hz), which has been widely associated with attentional engagement and cognitive processing. Alpha-band power estimates were extracted from the periodic component of the resulting model. For statistical analyses, alpha power was averaged across a posterior region of interest (ROI) identified in the grand-averaged scalp topography across all conditions. This cluster included 13 posterior electrodes. Mean alpha power values were then obtained for each participant and condition. Alpha power was computed separately for trials containing disruptions and trials without disruptions (Quiet condition). Differences between conditions were assessed using paired-sample t-tests across participants.

##### d) Phasic and Tonic skin-conductance

Last, we assessed whether disruptions affected ongoing skin-conductance activity at the full-trial level. Using the Ledalab decomposition output, we extracted trial-wise estimates of the **phasic driver** and **tonic activity** from the GSR signal. For each trial, the mean phasic and tonic signals were computed across the full trial duration, and then averaged separately for the Disruption condition and the Quiet condition for each participant. Differences between conditions were assessed using paired-sample t-tests.

### 3) Local-effects of disruptions – *during* and immediately *after* the disruption

Besides assessing the effects of disruptions at the full-trials level, we also analyzed neural and GSR data a more fine-grained temporal resolution, and assessed responses *during* and *after* disruption events, in five consecutive epochs. To this end, trials containing disruptions were further segmented into 2-second long epochs as follows (see illustration in Figure 1): *Epoch T* was defined from the onset of the disruption. *Epoch T+1* was defined from 1-second after the offset of the disruption, and Epochs *T+2, T+3, and T+4* followed consecutively with 0.1 s gaps between them. Note that since the inter-stimulus interval between disruptions was 10.75- 12.33s, epoch *T+4* immediately preceded the next disruption. Since the experiment contained 36 disruptions, this was the number of segments included in each epoch. In order to compare these responses to the Quiet condition, we randomly extracted 36 epochs from the Quiet condition to serve as controls, while ensuring similar sampling distribution over time as for the epochs in the Disruption condition.

**Figure 1.**
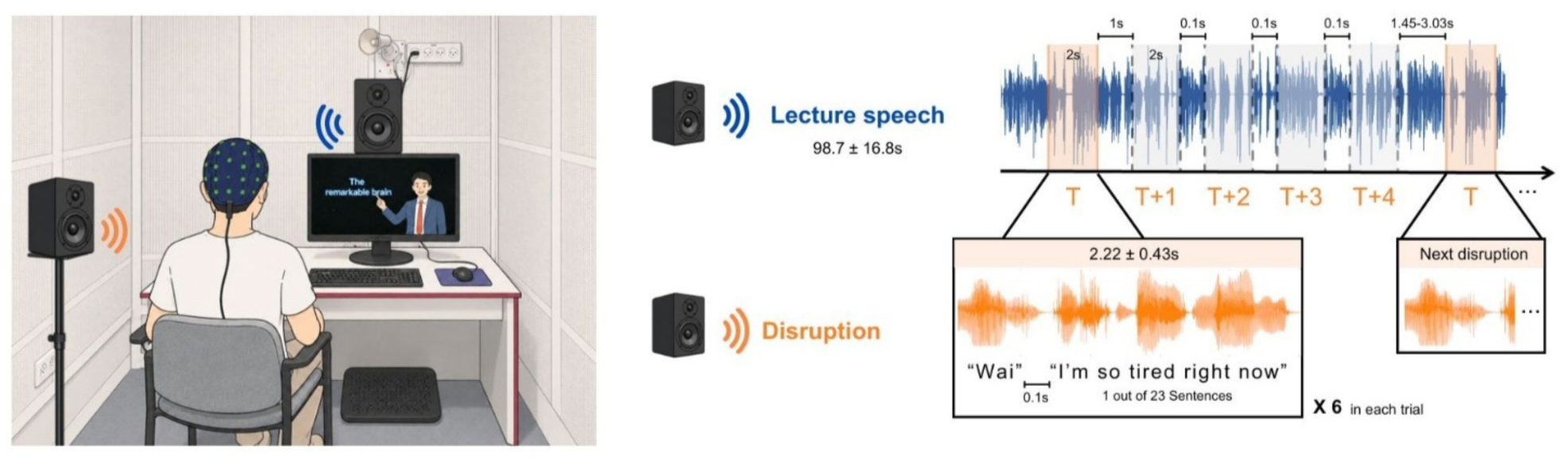
Schematic illustration of the time-resolved analysis. Trials containing Disruptions were segmented into five consecutive 2-s epochs time-locked to each disruption event, from disruption onset (T) through the post-disruption period (T+1–T+4).

For each epoch, we estimated the same four neural and skin-conductance metrics described above for the full-trial analyses, with slight adaptations to the shorter epochs, when needed:

*a) Speech tracking analysis:* Speech tracking analysis relies on training encoding/decoding models on large portions of data, and is therefore not easily applicable to short segments of speech (Mesik & Wojtczak, 2023). In order to assess speech-tracking responses in time-resolved 2-second epochs, we trained encoding/decoding TRF models on long segments of data taken from the Quiet condition and then tested them on the shorter epochs. Specifically, trials from the Quiet condition were segmented into 25-second long segments (3 segments per trial, separated by 4 seconds each). These data served as training data, whereas the left-out interspersed 4-second portions were used for testing (further divided into 2-second long epochs). This approach ensured similar sampling of the data for model training and tested across the experiment, while preserving sufficiently long segments of data for TRF training. Model training and λ optimization followed the same optimization procedure described above (“full trial analysis”). As in the trial-level analysis, model significance was assessed relative to a null distribution generated from permutation (100 permutation per participant). The resulting optimized model was then applied to epochs from the Disruption Condition (T, T+1, T+2, T+3, T+4) and the matched epochs from the Quiet condition (using the mTRFpredict.m function). We then assessed whether there were significant differences across epochs in model predicative power (encoding; averaged across the 40 electrodes identified in the full-trial analysis as showing significant speech tracking) or reconstruction accuracy (decoding) using a repeated-measures ANOVA followed by post-hoc pairwise comparisons.
*b) Inter subject correlation (ISC):* ISC was computed separately for each Epoch data from the Disruption Condition (T, T+1, T+2, T+3, T+4) and the matched epochs from the Quiet condition, using identical methods as describe above for the full-trial analysis (CCA decomposition followed by estimation of each individuals’ similarity to the group). We then assessed whether there were significant differences in ISC across epochs using a repeated-measures ANOVA, followed by Bonferroni-corrected post hoc pairwise comparisons.
*c) Alpha power Analysis:* Spectral estimates of alpha power were computed for each Epoch data from the Disruption Condition (T, T+1, T+2, T+3, T+4) and the matched epochs from the Quiet condition, using identical methods as describe above for the full-trial analysis, (calculation of PSD, followed by FOOOF correction). The mean alpha power was extracted from the posterior ROI of electrode for each participant and epoch. We then assessed whether there were significant differences in alpha power across epochs, using a repeated-measures ANOVA, followed by Bonferroni-corrected post hoc pairwise comparisons.

### Additional Exploratory Correlation Analyses

Given the large number of neurophysiological measures extracted here, we conducted exploratory correlation analyses to examine possible associations among the different measures, including: ERP component amplitudes (to disturbances), SCR peak amplitude, TRF encoding and decoding accuracy, mean ISC and alpha power. Additionally, we examined whether any of these neurophysiological responses, including local disturbance-related effects (i.e., the contrast between time-resolved epochs: T vs. Quiet or T vs. T+1) were associated with individual variability in ASRS scores, which capture self-reported symptoms of attentional difficulties. All correlations were computed using two-tailed Pearson correlations and FDR correction was applied separately within each analysis family.

## Results

### Behavioral results

To assess participants’ comprehension of the lecture content, we calculated accuracy rates on the multiple-choice questions presented after each segment. Overall, performance was high (M = 83.98%, SEM = 0.14%), confirming that participants successfully attended to and understood the material. Importantly, performance levels showed sufficient variability to rule out ceiling effects. No significant difference in accuracy was observed between Quiet trials and trials with Disruptions [Quiet (M = 84.90%, SEM=1.26%), Disruptions (M = 83.05%, SEM=1.46%, t(59) = 1.15, p = 0.256].

### Event-related responses to disruption onsets (“Wai”)

Both EEG and GSR signals showed clear transient responses elicited by the onset of disruptions (the interjection “Wai”; Figure 2).

**Figure 2.**
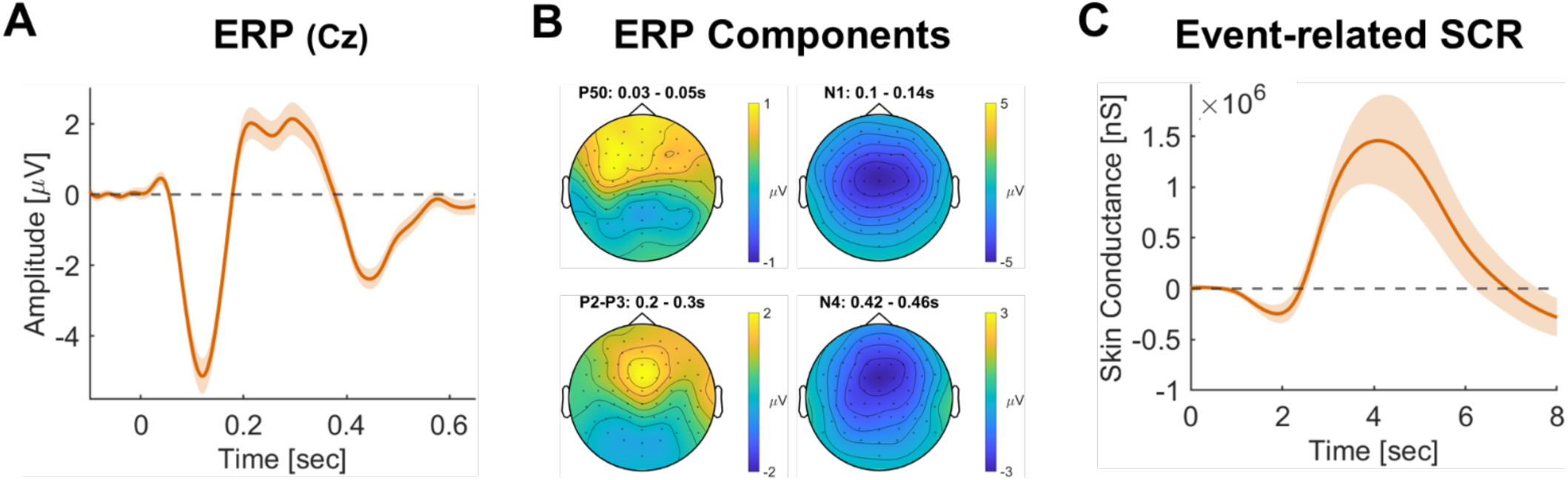
Neural and physiological responses to disruptions. **A.** Grand-averaged ERP at electrode Cz, time- locked to the onset of speech disruptions (n=60); shaded area indicates the SEM. **B.** Scalp topographies of four ERP components: P50 (30-50ms), N1 (100–140 ms), P2-P3 complex (200–300 ms) and N4 (420–460 ms). **C**. Grand-averaged event-related skin conductance response in nanoSiemens (n = 59), showing a phasic increase peaking around 4 s after onset; shaded area indicates SEM.

*EEG:* ERPs time-locked to disruption onset showed a typical response profile for auditory responses (Figure 2A). The grand-average waveform exhibited a sequence of components: an early P50 (30–50 ms), followed by an N1 (100–140 ms), a broad P2–P3 complex (200–300 ms), and a later N4 component (420–460 ms). As shown in Figure 2B, all responses displayed fronto-central scalp distributions.

*GSR:* The event-related skin conductance response (Event-Related SCR) also showed a clear phasic increase following disruption onset, peaking approximately 4 seconds after the event (Figure 2C), which is consistent with time time-course of autonomic responses. This response was significantly larger relative to time-matched segments extracted from the Quiet condition (t(58) = 5.10, p < .001).

### Effects of Disruptions on Speech tracking of Lecture

#### Global effects: Full-Trial analysis

Speech-tracking analysis applied at the full-trial level showed reliable speech tracking in a cluster of 40 fronto-central electrodes where the predictive power of the encoding model exceeded that of a permutation-derived null distribution (Figure 3A, inset). These electrodes were used for subsequent comparisons of TRF time-courses between conditions. The grand-averaged TRF showed the typical speech-evoked response profile with three prominent peaks: an early positive peak at 30–50 ms (corresponding to the P50 component), followed by a negative deflection at 80–100 ms (N1), and a later positive peak at 160–180 ms (P2) (Figure 3A). While the overall TRF morphology was similar across conditions, the magnitude of the response differed between trials with and without disruptions. Statistical comparisons showed no significant difference in the P50 window (t(59) = −0.96, p = .160), but both the N1 and P2 responses were significantly larger in Quiet vs. Disruption trials [t(59) = 2.62, p = .008 and t(59) = −2.95, p = .006, respectively]. Topographical maps of these effects showed fronto-central distributions consistent with canonical speech-tracking responses (Figure 3B). Complementary decoding analyses showed a similar trend, with higher speech reconstruction accuracy in Quiet vs. Disruption trials, although this difference was only marginally significant [t(59) = -1.7, p = 0.094; Figure 3C].

**Figure 3.**
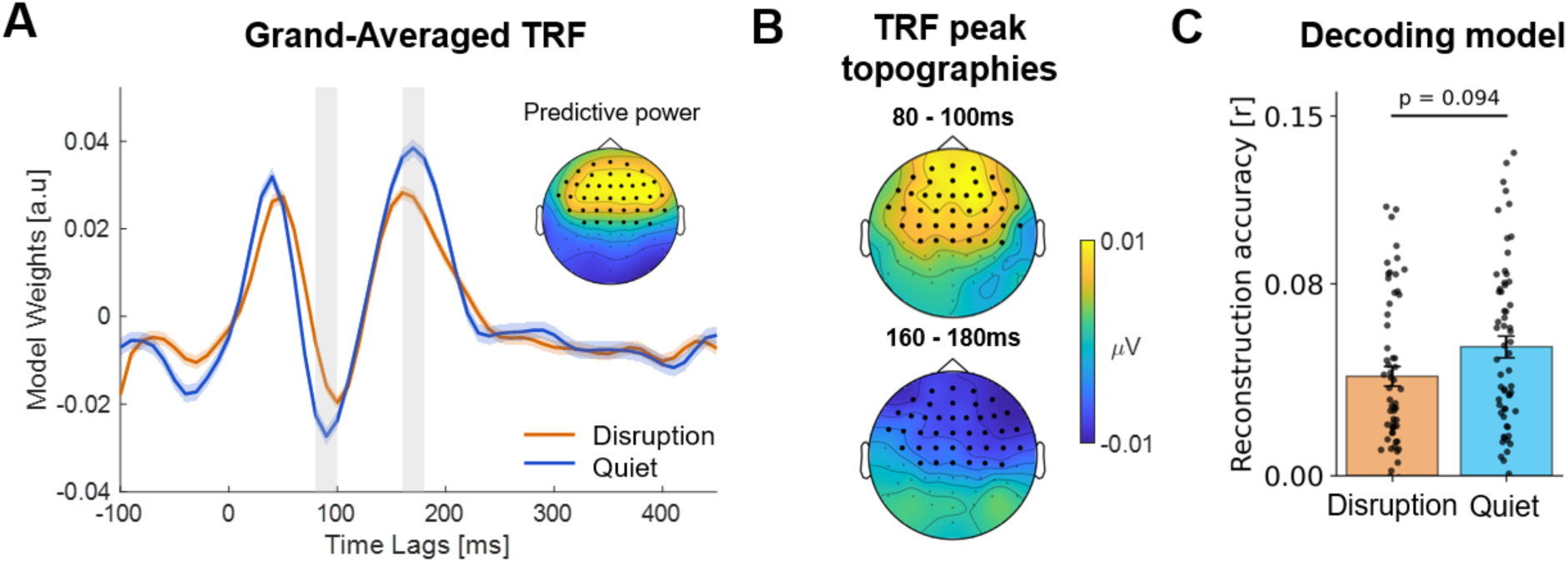
Global effects of disruptions on neural tracking of the lecture speech (full-trial analysis). **A.** Grand-averaged encoding TRF weights for the Quiet (blue) and Disruption (orange) conditions (orange). Shaded regions mark windows with significant differences between conditions (80–100 ms and 160–180 ms). The inset shows the scalp distribution of encoding predictive power; black dots indicate the 40 electrodes exceeding the permutation-derived null distribution and used for subsequent analyses. **B.** Topographical maps of condition differences in TRF weights at the N1 and P2 latency windows, with black dots indicating electrodes showing significant condition differences. **C.** Decoding results showing reconstruction accuracy of the speech envelope the Quiet and Disruption conditions. Dots represent individual participants and bars indicate the group mean.

### Local-effects of disruptions: Time-resolved analysis

We next examined the effect of disruptions on speech tracking of the lecture at a more temporally resolved level, to assess whether the effects observed at the global-level stem from the portions of the trial with overlapping audio (epoch T) or if effects continued post-disruption (epochs T+1 to T+4).

First, we confirmed that the TRF estimated for the time-resolved analysis was similar to that estimated in the full-trial analysis and contained the same three canonical peaks (P50, N1, P2; Figure 4A). We then assessed how the predicative power of speech tracking models varied across epochs. Encoding results (Figure 4B, top) revealed a significant main effect of Epoch [F(5, 295) = 7.94, p < .001], with post hoc comparisons indicating that speech-tracking during Epoch T (overlap with disruption) was significantly lower relative to the Quiet condition and relative to all post-disruption epochs (T+1 to T+4; all p < .01, except T vs.T+3 with p=0.045). In contrast, no difference was found between speech-tracking during the Quiet condition vs. any of the post-disruption epochs (T+1 through T+4; all ps > 0.05), which also did not differ among themselves [all ps = 1.000]. A similar pattern was observed also for the decoding analysis (Figure 4B, bottom), with a significant main effect of Epoch [F(5, 295) = 11.45, p < .001], stemming from lower reconstruction accuracy during Epoch T relative to all other epochs (all p < .001), and no other significant differences between epochs (all p = 1.000). Together, these results suggest that the differences observed at the full-trial level were primarily driven by reduced speech tracking in the epoch when disruptions were presented, but did not carry-over to post-disruption epochs (T+1 to T+4), which did not differ from responses in the Quiet condition.

**Figure 4.**
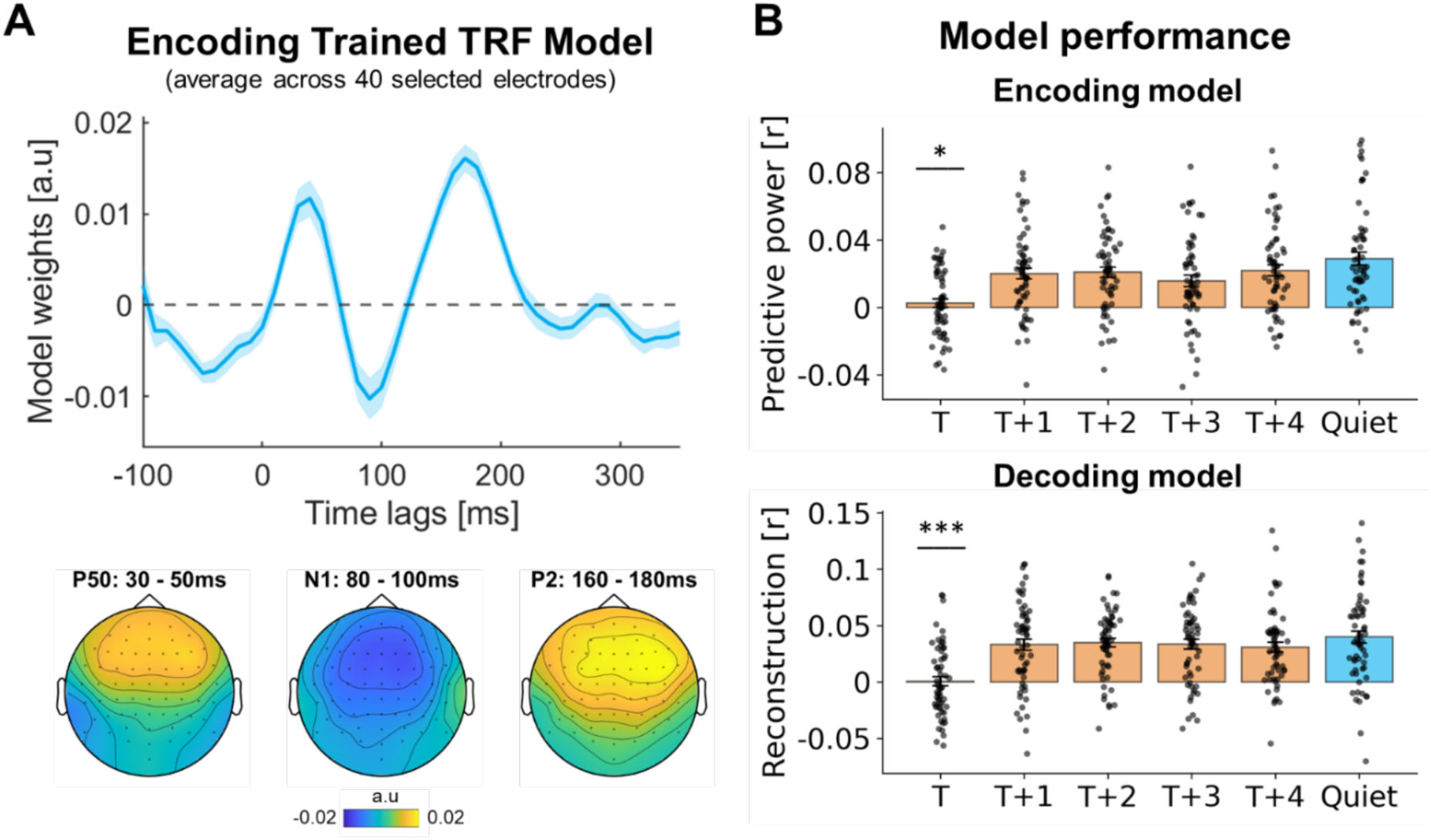
Time-resolved effects of disruptions on neural tracking of the lecture speech. **A.** Grand-averaged encoding TRF weights estimated for the time-resolved analysis (averaged across 40 electrodes selected in the full-trial analysis). The temporal response profile (top) and scalp topographies (bottom) are in line with canonical auditory components (P50: 30-50ms, N1: 80-100ms, P2: 1680-180ms). Shaded regions indicating the SEM. **B.** Speech tracking predictive power (encoding model; top) and reconstruction accuracy (decoding model; bottom) in time-resolved epochs T to T+4 around disruptions (orange) and matched epochs from the Quiet condition (blue). Bars represent the group mean (Pearson’s r), dots indicate individual participants, and error bars denote SEM. Asterisks indicate significant differences between the disruption epoch (T) and all other epochs (* p < .05, *** p < .001).

### Inter subject correlation (ISC)

To further examine shared neural dynamics, we quantified inter-subject correlation (ISC) using CCA-based components. ISC was calculated based on the average of the first three components, that reflect the spatial patterns capturing the most shared variance across participants (Figure 5A). When analyzed at the *full-trial level*, mean ISC was slightly higher in the Disruption condition than the Quiet condition, although this difference was marginally significant [t(59) = 1.8, p = .076; Figure 5B]. When analyzed in a *time-resolved* manner we found a significant main effect of Epoch [F(5, 295) = 112.849, p < .001; Figure 5C]. Post hoc comparisons indicated that ISC values during the epoch containing disruptions (Epoch T) were significantly higher relative to all other epochs, including the Quiet epoch (all p < .001), whereas none of the other epochs differed from each other (all p > .05). Together, these results suggest that the trend observed in the full-trial ISC analysis was primarily driven by increased ISC in the epoch where disruptions were presented, but did not carry-over to post-disruption epochs (T+1 to T+4).

**Figure 5.**
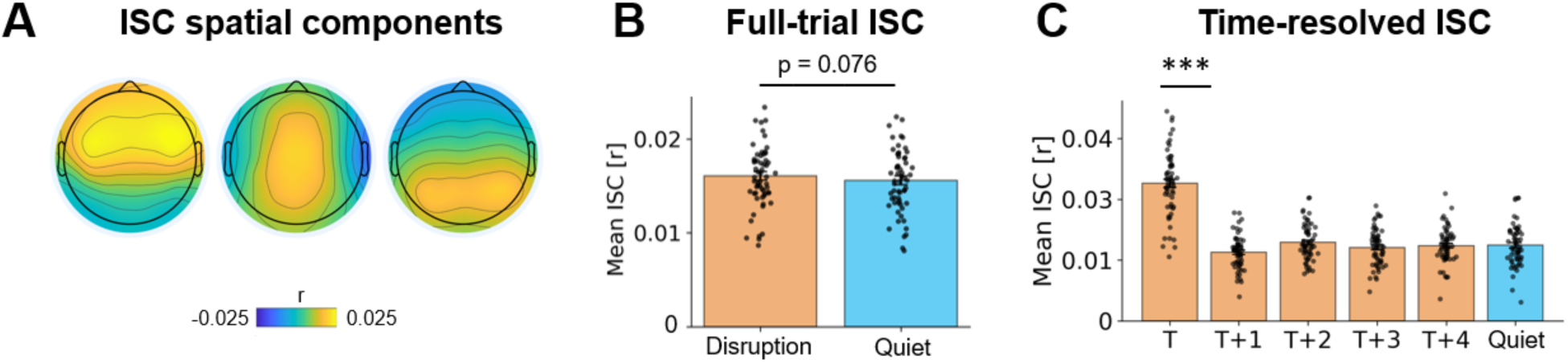
Inter-subject correlation (ISC) during lecture listening. **A.** Spatial maps of the first three correlated components identified by CorrCA, capturing shared variance in neural activity across participants. **B.** Mean ISC values calculated at the full-trial level in the Quiet (blue) and Disruption (orange) conditions. Bars indicate the group mean and dots represent individual participants. **C.** ISC values calculated in a time-resolved manner in the disruption condition (Epochs T–T+4, orange) and matched Quiet epoch (blue). Bars and dots represent the group means and individual participants as in B. Asterisks (***) indicate significant differences between Epoch T and all other epochs (p < .001).

### Alpha-power analysis

The grand-averaged power spectrum revealed a clear alpha peak, which defined the 7–13 Hz band used for subsequent analyses (Figure 6A). The corresponding scalp topography showed a posterior distribution, and the posterior ROI used for statistical analysis is indicated by black dots (Figure 6A, inset).

**Figure 6.**
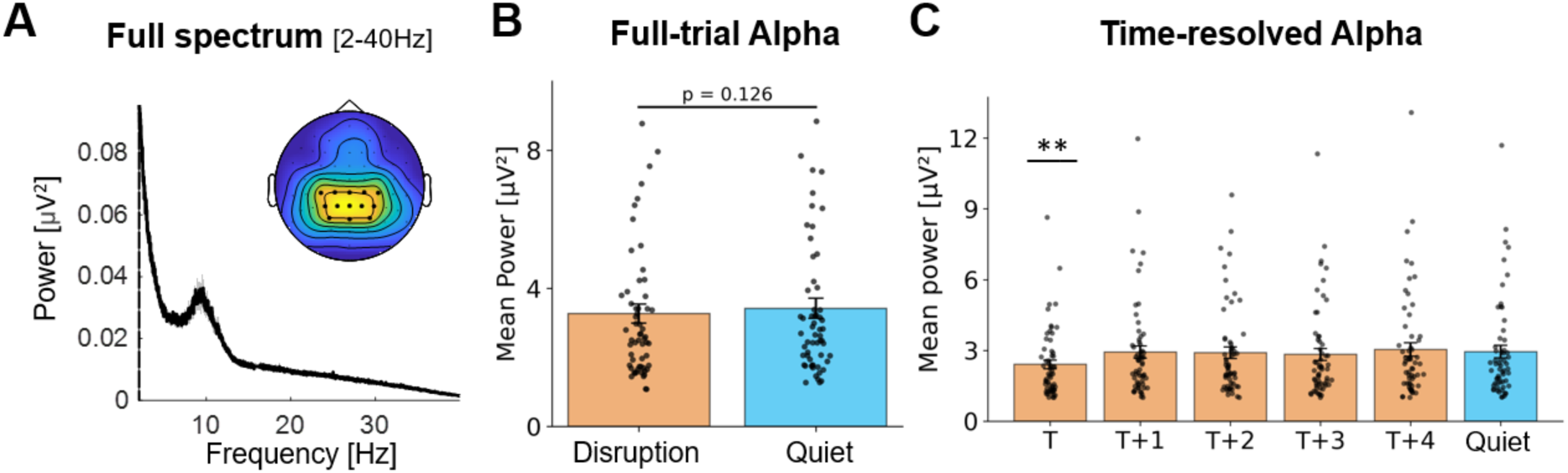
Alpha-band power during lecture listening. **A.** Grand-averaged power spectrum (2–40 Hz) showing a clear alpha peak, and corresponding scalp topography (7–13 Hz); black dots indicate the posterior ROI used for analysis. **B.** Mean alpha power in trials with disruptions (orange) and Quiet trials (blue). Bars indicate the group mean and dots represent individual participants. **C.** Alpha power calculated in a time-resolved manner in the disruption condition (Epochs T–T+4, orange) and matched Quiet epoch (blue). Bars and dots represent the group means and individual participants as in B. Asterisks (**) indicate significant differences between the epoch T and all other epochs (p < .01).

When analyzed at the *full-trial level,* mean alpha power was slightly lower in the Disruption condition than in the Quiet condition, although this difference did not reach statistical significance (t(59) = -1.55, p = .126; Figure 6B). When analyzed in a *time-resolved* manner we found a significant main effect of Epoch [F(5, 295) = 8.08, p < .001; Figure 6C]. Post hoc comparisons indicated that alpha power was significantly reduced during the epoch containing disruptions (Epoch T) compared to the matched Quiet epoch and all post-disruption epochs (T+1 to T+4; all p ≤ .002). In contrast, Alpha power in the post-disruption epochs did not differ from each other or from the Quiet epoch (all ps = 1.000). Together, these findings suggest that the pattern observed in the full-trial analysis was primarily accounted for by reduced alpha power during the disruption epoch (T), rather than by differences across the remaining epochs.

### Phasic and tonic skin-conductance

To assess whether disruptions affected ongoing autonomic activity, we analyzed the phasic and tonic skin-conductance activity. When analyzed at the *full-trial level*, neither tonic or phasic activity were significantly different in the Disruption condition than the Quiet condition [t(58) = -0.07 ,p = .94; t(58) = 0.14, p = .89 respectively, Figure 7). This analysis could not be reliably applied in a more *time-resolved* manner due to the slow nature of the skin-conductance response (Benedek & Kaernbach, 2010a; Cacioppo et al., 2007).

**Figure 7.**
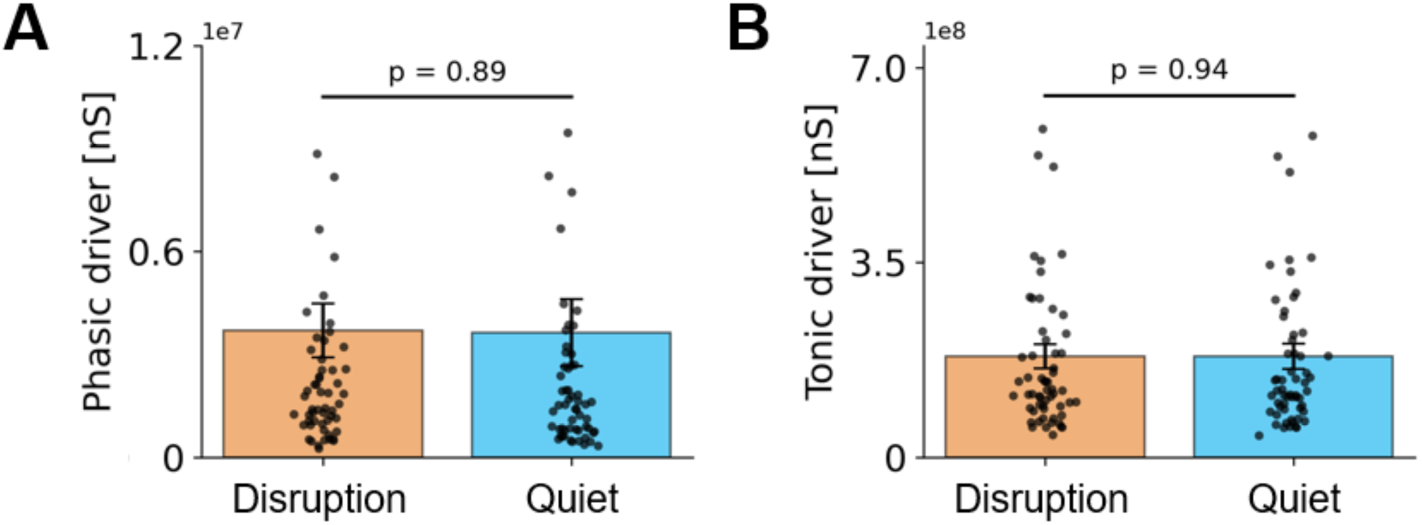
Skin-conductance activity during lecture listening. A. Mean phasic skin-conductance activity for trials with disruptions (orange) and quiet trials (blue). B. Mean tonic skin-conductance activity for trials with disruptions (orange) and quiet trials (blue). Dots represent individual participants and bars indicate the group mean.

### Exploratory correlation analyses

We conducted an exploratory analysis of the correlations among the different neurophysiological measures during the disturbance window itself (T) and the amplitude of the event-related neural and SCR responses to the disturbances. As shown in Table 1, the overall pattern of associations was sparse, with most correlations being small and non-significant. At the uncorrected level, mean ISC was positively correlated with alpha power (r = -.29, p = 0.02), ERP N1 (r = -.34, p = 0.008), ERP P2-3 (r = .38, p = 0.003), and ERP N4 (r = -.27, p = 0.04) amplitudes, and ERP P50 was positively correlated with ERP P2-3 amplitude (r = .35, p = 0.006). After FDR correction across the unique pairwise correlations, only the association between ERP N1 and ERP N4 amplitudes remained significant (r = .49, p < 0.001, pFDR = .002). This sparse pattern suggests that the different neurophysiological measures captured partly independent facets of the response to auditory disturbances, rather than reflecting a single shared neural response.

**Table 1.** Correlations among neurophysiological measures. Pairwise correlations between the various neural measures extracted during epoch with disruptions (T; variables 1-4) and event- related neural and SCR responses to disruptions (variables 5-9). Values indicate Pearson’s r. † indicates p < .05 uncorrected. * indicates p < .05 after FDR correction.

| Variable | 1 | 2 | 3 | 4 | 5 | 6 | 7 | 8 | 9 |
| --- | --- | --- | --- | --- | --- | --- | --- | --- | --- |
| 1. Alpha power (T) | - |  |  |  |  |  |  |  |  |
| 2. Mean ISC (T) | -.29<sup>†</sup> | -- |  |  |  |  |  |  |  |
| 3. TRF Encoding (T) | -.06 | .12 | - |  |  |  |  |  |  |
| 4. TRF Decoding (T) | -.01 | .05 | .19 | - |  |  |  |  |  |
| 5. SCR | -.03 | -.02 | .03 | .04 | - |  |  |  |  |
| 6. ERP P50 | .00 | .03 | .08 | -.23 | -.00 | - |  |  |  |
| 7. ERP N1 | -.04 | -.34<sup>†</sup> | .00 | .09 | -.18 | -.19 | - |  |  |
| 8. ERP P2-3 | -.08 | .38<sup>†</sup> | .20 | .11 | .15 | .35<sup>†</sup> | -.10 | - |  |
| 9. ERP N4 | -.04 | -.27<sup>†</sup> | -.03 | .12 | .07 | -.17 | .49<sup>*</sup> | -.14 | - |

In a second exploratory analysis, we tested whether the effects of disruptions on different neurophysiological measures were correlated with ASRS scores (Figure 8), which captures introspective self-reported of difficulties in attention in real life. We focused on response to the disruptions themselves (as in Table 1) as well as on the effect of disruptions relative to quiet (T vs. Quiet) and the return to baseline following disruptions (T vs. T+1). The correlation coefficients of all measures are shown in Table 2, however none of them were statistically significance (all pFDR > 0.05).

**Figure 8.**
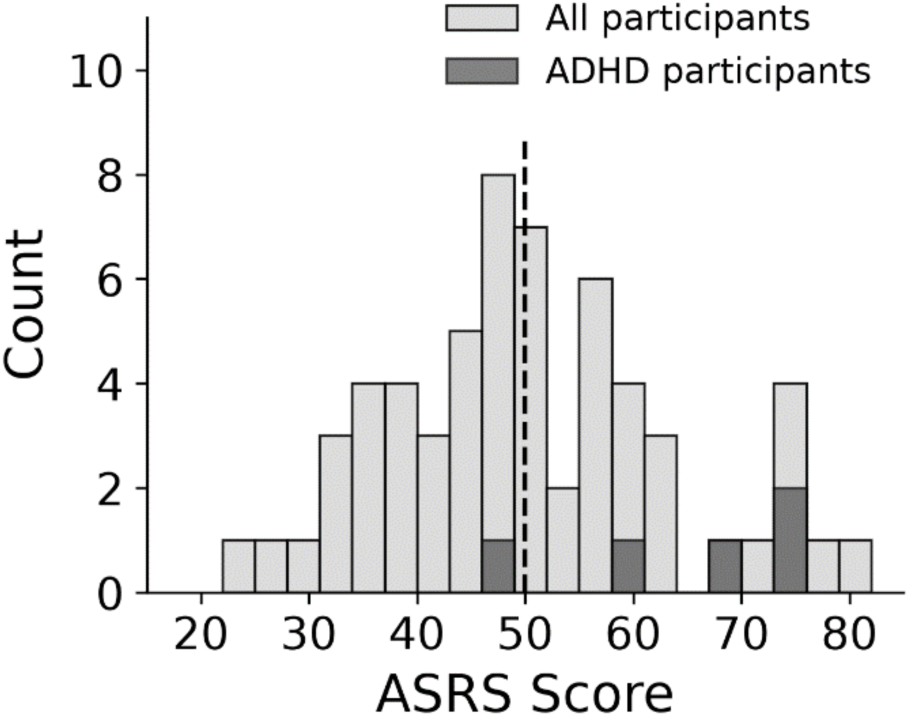
Distribution of ASRS scores across participants. Light gray bars represent the full sample, whereas dark gray bars indicate participants with a prior ADHD diagnosis. ASRS scores showed substantial variability, with many participants scoring above 51 (predefined ASRS screening threshold used to indicate elevated ADHD symptom levels; indicated by the vertical dashed line).

**Table 2.** Correlations between ASRS scores and neurophysiological responses to disruptions. Values indicate Pearson’s r of ASRS scores vs. each of the neurophysiological measures, shown in separate columns for responses to the disruptions themselves and for the effects of disruptions relative to quiet (T vs. Quiet) and the return to baseline following disruptions (T vs. T+1). † indicates an uncorrected p < .05; none of the correlations were significant after fdr correction for multiple comparisons.

| Measure | Correlations with<br>response to disruptions | Correlations with<br>T vs. Quiet contrast | Correlations with<br>T vs. T+1 contrast |
| --- | --- | --- | --- |
| Alpha power (T) | -0.20 | -0.01 | 0.06 |
| Mean ISC (T) | -0.06 | 0.17 | -0.20 |
| TRF Encoding (T) | 0 | 0.17 | -0.21 |
| TRF Decoding (T) | 0.08 | 0.05 | -0.06 |
| Event-Related SCR | 0.15 |  |  |
| ERP P50 | -0.12 |  |  |
| ERP N1 | 0.17 |  |  |
| ERP P2-3 | -0.02 |  |  |
| ERP N4 | 0.26<sup>†</sup> |  |  |

## Discussion

The present study examined the transient and global impact of background speech- disruptions on neurophysiological measures and comprehension during video-based learning. We found that disruptions elicited robust neurophysiological responses, capture both in a stimulus-evoked neural response and phasic increase in skin conductance. Disruptions were also associated with a **cost** to neural speech processing of the lecture, reflected in reduced neural tracking, as well as increased inter-subject correlation and decreased alpha power. Interestingly, these effects were **transient in nature,** limited only to the time-period in which disruptive speech was presented, whereas once the disruption ended, neural responses returned to levels comparable to the Quiet condition. These findings suggest that although background speech can be disruptive during tasks that require continuous attentional focus, due both to perceptual masking (Brungart, 2001; Mattys et al., 2012; Wang & Xu, 2021) as well as attentional capture (Bell et al., 2022; Marsh & Jones, 2010), nonetheless the auditory and/or attention systems are adept at recovering from bouts of disruption and re-engaging with target speech, with no apparent long-lasting disruptive effects.

### Neurophysiological Responses to Speech Disruptions

In contrast to classic auditory-attention paradigms, where disruptive stimuli typically consist of simple and semantically-meaningless sounds, here disruptions consisted of meaningful and contextually relevant speech utterances that carry both acoustic salience as well as semantic content. These disruptions elicited clear neural and physiological responses, in line with bottom-up “orienting”. Neural response manifest in a classic-looking auditory ERP sequence, including an early positive P50 response (30-50 ms), a negative N1 (100-140 ms), a broad positive complex akin to a P2/P3 response (200-300 ms), and a later negative component (420-460 ms). This temporal profile naturally invites comparison with the classical auditory-distraction literature, where rare deviant sounds (e.g., in oddball paradigms) elicit a stereotyped "distraction potential", such as the early mismatch negativity (MMN; ∼150-250 ms) that is related to automatic change detection and the P3a (∼250-350 ms) that is associated with involuntary attentional orienting toward the deviant. However, applying these canonical labels, that are derived from highly controlled event-related designs, to data collected under more continuous and naturally-uncontrolled contexts is not straightforward (Fiedler et al., 2025; Justen & Herbert, 2018; Korte et al., 2025; Levy, Libman Hackmon, et al., 2025; Straetmans et al., 2022). Therefore, while noting the parallels between the current results and more traditional attention studies, we must also bear in mind the substantial difference between analyzing responses to short transient events vs. continuous, ecological speech (Bell et al., 2010; Rosenkranz et al., 2024).

And yet, in the present paradigm disturbances were designed to preserve event-related features. Specifically, since each disruption began with the same interjection “Wai”, this provided the temporal alignment necessary to generate an ERP (despite being presenting on the background of continuous speech). Accordingly, we feel confident in interpreting the sequence of ERP components to the “Wai” disturbance in relation to the traditional auditory ERP literature, namely that the early P50 and N1 reflect sensory registration and detection of the event, whereas the broad P2/P3 complex parallels the P3a elicited by novel yet task-irrelevant sounds, which has been linked to attentional orienting, stimulus evaluation, and context updating despite their behavioral irrelevance (Escera & Corral, 2007; Friedman et al., 2001; Polich, 2007). Regarding the later negative response observed around 420–460 ms, we can offer two possible interpretations. One is that it is type of N400 response, found in response to semantically-surprising words (Kutas & Federmeier, 2011; Kutas & Hillyard, 1980). Although N400 responses are sometimes attenuated for words that are actively ignored (Erlbeck et al., 2014), (Brown et al., 2023) showed that presenting background speech with semantically-salient words in a realistic environment, such as ones’ name, elicit both P300 and N400-like responses. Accordingly, one interpretation is that the late negative response observed here is a reflection of semantic analysis applied to the distractor speech. Another possibility is that this response is similar to the Reorienting Negativity (RON) that has been linked to re-engagement with a primary task following disruptions (Escera & Corral, 2007; Näätänen et al., 1978; Schröger & Wolff, 1998; Wetzel et al., 2004). According to this interpretation, the late negatively may reflect active efforts to withstand distraction by the disturbance and maintain attention towards the target speech. As additional research is accrued on the continuum between event-related and continuous speech stimuli, we hope to gain a clearer understanding of this late response and the relationship between traditional ERP components and those generated under more ecological conditions.

Complementing the ERP findings, measurement of **skin conductance** shows a strong, phasic autonomic arousal response to disruptions. This response, sometimes referred to as an “orienting response”, is typically observed following unexpected yet salient events (Barry et al., 2023; BRADLEY, 2009; Critchley, 2002; Zimmer & Richter, 2023), as well as stimuli with personal relevance or social/emotional value (Brown et al., 2023; Pinto et al., 2023) (Aue et al., 2011; Carbone et al., 2025; Dindo & Fowles, 2008; Gronau et al., 2003). The slow temporal dynamics of this response (SCR peaks after ∼4 seconds; Benedek and Kaernbach 2010; Cacioppo, Tassinary, and Berntson 2007) prohibits us from establishing whether it is driven primarily by acoustic salience or also by its semantic features. Nonetheless, the SCR orienting response complements the ERP responses to indicate that background disturbances elicit potent bottom-up responses. That these responses emerged despite active engagement in watching the lecture underscores the difficulty of sustaining selective attention in naturalistic listening environments, and sets the stage for examining how this orienting shaped the neural processing of the lecture itself.

### The “Cost” of Disruption

Beyond eliciting neurophysiological orienting responses, disruptions also effected other features of the EEG. Specifically, disruptions were associated with neural tracking of the teacher’s speech (TRF), well as an increase in inter-subject correlation (ISC) and decrease in alpha power. While these effects were modestly significant when analyzing results at the global trials-wise level, the more time-resolved analysis revealed they were primarily driven from the 3-second period during which the disruptions were presented and overlapped with the lecture itself. In other words, these effects were transient rather than long-lasting.

The observed reduction in speech tracking is consistent with the well-established finding that cortical tracking speech is modulated by selective attention (Fiedler et al., 2019; Har-Shai Yahav et al., 2024; Schüller et al., 2023; Yahav et al., 2025) and is diminished when attentional resources are diverted (Fuglsang et al., 2020; Ren et al., 2026; Vanthornhout et al., 2019; Zion Golumbic et al., 2013). Hence, the transient reduction in speech tracking may reflect a temporary diversion of processing resources away from the target speech, manifest either as an attention shift toward the disruptive speech (Broadbent, 1958; Treisman, 1964) or a division of resources among the two stimuli (Ding & Simon, 2012; Kaufman & Zion Golumbic, 2023). Given that in the current study disruptive speech was intelligible and semantically meaningful, target and disturbances likely compete for processing resources both the acoustic and linguistic levels (Dai et al., 2017; J. Song et al., 2020).

Disruptions were also accompanied by a transient increase in ISC and reduction in alpha- power. In interpreting these effects, we must note that – in contrast to the speech tracking response that is directly linked to the teacher’s speech – these metrics are not stimulus-specific but reflect more general features of the neural response. Specifically, ISC quantifies the degree of similarity in the neural activity across participants and is thought to reflect their “joint attention” or “level of engagement” with the presented stimuli (Cohen et al., 2018; Dmochowski et al., 2012; Ki et al., 2016; Rosenkranz et al., 2021). The finding that ISC increased when disruptions were presented indicates that the neural responses generated by these stimuli (e.g., the ERP discussed above) led to higher similarity across participants, relative to when they were “just” listening to the speech. Similarly, the decrease in alpha power can also be interpreted as reflecting a momentary increase in engagement following disruptions. Alpha power is famously known to decrease after salient events, a pattern attributed to transitioning from a baseline state into an active-processing mode (Klimesch et al., 2007; Pfurtscheller & Lopes da Silva, 1999; Woodman et al., 2022). Alpha oscillations have been hypothesized to play an important role in attention functioning, with lower alpha power generally associated with higher levels of focus (Foxe & Snyder, 2011; Jensen & Mazaheri, 2010; Klimesch, 1999) and selective enhancement of alpha in specific brain regions associated with inhibition of stimuli in irrelevant locations or modalities (Jensen & Mazaheri, 2010; Kelly et al., 2006; Worden et al., 2000). Here, we attribute the decrease in alpha following disruptions as another feature of the orienting response towards them, involving both a momentary breakdown of the internally maintained state that supports ongoing lecture processing, and a transition toward a more externally driven, stimulus-reactive mode. This aligns with evidence linking sustained alpha to working memory maintenance and internally directed processing (Cooper et al., 2003; Poch et al., 2014; Xie et al., 2016), as well as findings showing that involuntary orienting to distractors is accompanied by alpha desynchronization (Arana et al., 2022; Fodor et al., 2020; Harris et al., 2017). Importantly, only modest and partial correlations among the different neural manifestations of the response to disruptions (ERPs, reduction in speech tracking, change in ISC and alpha power) which suggests that they largely reflect independent mechanistic facets of how the brain deals with disruptions, rather than neural epiphenomena.

### Post-Disruption Re-engagement

A central question posed in this study is whether the occurrence of disruptions has global ramifications for task performance and target speech processing, beyond the time of the disruption itself. By conducting a time-resolved analysis of neural responses in time-windows during and immediately following disruption offset we were able to investigate the time-course for “recovery” from disruption. Although one could hypothesize that the presence of disruptions would reduce attention to the lecture more globally, this was not the case here. We found that all three neural measures that changed during the disruption themselves (speech tracking, ISC and alpha power) returned to baseline levels in the epoch immediately following disruption offset (T+1), with no statistically significant differences between post-disruption and quiet epochs at any subsequent time point (T+1 through T+4). These results suggest that the neural cost of the disruptions was limited to the disruption window, with no evidence for prolonged negative effects. Rather, these findings are consistent with accounts describing swift re-orientating back to the primary task following involuntary capture by irrelevant stimuli (Escera et al., 2001; Schröger & Wolff, 1998). Similarly, we did not find any differences in performance between trials with and without disruptions, which further supports the time-limited effect of disturbances since – given their short duration - their acoustic masking alone would not necessarily lead to a substantial loss of information that would impair overall comprehension. Overall, the pattern observed here aligns with the view that the auditory attention system operates with considerable flexibility and resilience, and is adept at keeping focus on a target stream despite occasional disruptions.

This attentional resilience is in line with the notion that auditory attention is supported by maintaining a structured multi-dimensional representation of the target stream. According to prevalent theories of Auditory scene analysis, features belonging to the same source are bound by shared temporal, spatial, spectral and object-related regularities, which helps maintain expectations about target speech (Bendixen, 2014; Bregman, 1990; Shamma et al., 2011; Winkler et al., 2009). Within this framework, brief disturbances do not fully destabilize the multi-dimensional representations of target speech, which remains robust during disturbances and supports rapid re-engagement once disturbances end.

Although in the current study the disruption-related effects observed here were temporally bounded with no lingering cost, other studies have found that task-irrelevant sounds can sometimes have longer-lasting consequences. For example, Straetmans et al. (2022) showed reduced neural tracking of both target and non-target speech streams in a 5- s window following a salient environmental sounds, and Holtze et al. (2021) actually found that neural tracking of target speech increased after hearing one’s own name. Studies monitoring pupil dilation and blinks also report that distractor-related arousal can persist into subsequent target processing (Fiedler et al., 2025; Liu & Chait, 2026). But even in those studies, the effects of disturbances are relatively short-lived and are followed by a return to baseline. Integrating across studies, these transient effects are consistent with a push-pull dynamic described in a previous study by Huang & Elhilali (2020), indicating that local disruptions lead to a momentary shift in resources away from a main target stimulus, but do not necessarily have long-lasting detrimental effects.

### Relation to ADHD

Given that recovery from disruption may nonetheless vary across individuals, we examined whether this variability was related to self-reported ADHD symptomatology. ADHD is commonly associated with increased distractibility, reduced sustained attention, and greater difficulty maintaining task focus in naturalistic settings (Kofler et al., 2008; Lauth et al., 2006; Stokes et al., 2022; Tucha et al., 2017; Vile Junod et al., 2006). Accordingly, we could have expected that individuals with higher ASRS scores, who report experiencing more difficulties in attention in their daily life, would also exhibit more pronounced neurophysiological responses to disruptions or prolonged post-disruption recovery. However, this was not the case, as ASRS scores were not significantly correlated with any of the neurophysiological measures. This null pattern is sobering, but is also in line with recent meta-analyses describing the difficulties and inconsistencies in relating ADHD symptoms to specific objective neural-markers (Barkley, 2019; Faraone et al., 2021; Arrondo et al., 2023; Schweitzer and Zion Golumbic, 2023; Sonuga-Barke et al., 2023). Many factor contribute to these challenges, from the substantial heterogeneity of attention-related difficulties, through their high correlation with other social-emotion difficulties, as well as the reliance on self-reports which are highly subjective and influenced by individuals’ perspective on their own behavior (Butzbach et al., 2021). The current null results contribute to this ongoing conversation, and highlight a discrepancy between participants’ subjective evaluated their attention abilities ”in general”, as captured through their ASRS scores, and the plethora of objective behavioral and neurophysiological measures collected in this rather naturalistic task (Fuermaier et al., 2015; Kallweit et al., 2020; Toplak et al., 2013). Given increased concern about rising rates of attention-difficulties (Davidovitch et al., 2017; Danielson et al., 2024), we see these results as quite encouraging, as they suggest that most individuals have a good ability “bounce back” after disruptions and resume paying attention to the task at hand (Esterman et al., 2013, 2014 Esterman & Rothlein, 2019; H. Song et al., 2021, Körner et al., 2017).

### Methodological Considerations: Global and Local Effects

A methodological point that emerges from the present study concerns the value of combining global and local analyses. The global contrasts, which compared neural measures averaged across entire disruption trials with quiet trials, provided a coarse index of the overall disruption effect. However, because these measures average across the full trial, they are limited in their ability to distinguish sustained changes in processing from brief, time-locked disruptions. The local zoom-in analyses complemented this view by tracking how the effect changed over time, showing that the disruption-related changes were limited to the overlap window and resolved shortly after the disruptive speech ended. This is especially important for naturalistic listening paradigms, where attention and speech processing fluctuate over time rather than remaining stable throughout a trial. Global averages may therefore mask short-lived effects or mix together periods of disruption and recovery. Conversely, local analyses can reveal when an effect emerges, how long it lasts, and whether it carries over into subsequent processing. Future work would benefit from developing approaches that allow even finer temporal tracking of such fluctuations within naturalistic tasks. At the same time, this remains methodologically challenging, because smaller time windows provide less data and may reduce the reliability of estimates such as TRF, ISC, or spectral power (Mesik & Wojtczak, 2023; Nastase et al., 2019).

### Limitations and Future Directions

While this study represents a step forward in terms of ecological validity of attention research, it is still far from emulating the true challenges of paying attention to speech and dealing with disturbances in real-life environments. This study was still conducted in a quiet laboratory environment, and used recoded stimuli presented through loudspeakers, rather than generated by live people. In the future, we hope to extend this line of investigation into actual live environments, with multiple sources of perceptual salient and socially relevant disruptions, and to examine the effects of disruption and recovery dynamics under those conditions (Bronkhorst, 2015; Janssen et al., 2021; Ladouce et al., 2017; Röer & Cowan, 2021). Another limitation of the current experiment is the use of single lecture, whose content and delivery may have been more or less engaging to different participants, a factor that likely interacts with the treatment of disruptions (Levy, Shadi, et al., 2025). Hence, a more comprehensive study would require exploring how features of target speech and of disruptions interact to affect allocation of attention and competition for processing resources between the competing stimuli.

## Acknowledgements

This work was funded by the Israel Science Foundation (ISF), grant # 274/24, by the Binational Science Foundation (BSF) grant # 2022024 and by the National Institute of Mental Health (NIMH) grant # 5R01MH135266.

